# Is the production of reactive oxygen and nitrogen species by macrophages associated with better infectious control in the experimental disseminated and pulmonary mucormycosis?

**DOI:** 10.1101/2022.06.06.494943

**Authors:** Amanda Ribeiro dos Santos, Thais Fernanda Fraga-Silva, Débora de Fátima Almeida Donanzam, Angela Carolina Finatto, Camila Marchetti, Maria Izilda Andrade, Olavo Speranza de Arruda, Maria Sueli Parreira de Arruda, James Venturini

## Abstract

Different levels of resistance against *Rhizopus oryzae* infection have been observed between inbred (BALB/c) and outbred (Swiss) mice, with is associated with the genetic background of each mouse strain. Considering that macrophages play an important role in host resistance to *Rhizopus* species, we use the different infectious outcomes observed in experimental mucormycosis to identify the most efficient macrophages responses pattern against *R. oryzae in vitro* and *in vivo*. For this, we compared BALB/c and Swiss macrophage activity pre-and-post intravenous or intratracheal *R. oryzae* infections. Production of hydrogen peroxide (H_2_O_2_) and nitric oxide (NO) was determined in cultures of peritoneal (PMΦ) or alveolar macrophages (AMΦ) challenged, or not, with heat-killed spores of *R. oryzae*. Levels of TNF-α and IL-10 were also measured to enhance our findings. Naïve PMΦ from BALB/c increased the production of H_2_O_2_, TNF-α, and IL-10 in the presence of heat-killed spores of *R. oryzae*, while naïve PMΦ from Swiss mice was less responsive. Naïve AMΦ from two strains of mice were less reactive to heat-killed spores of *R. oryzae* than PMΦ. On 30 days of *R. oryzae* intravenous infection, lower fungal load in BALB/c strain of mice was accompanied by higher production of H_2_O_2_ by PMΦ when compared with Swiss mice. Differently, AMΦ from BALB/c mice showed higher production of NO, TNF-α, and IL-10 after 7 days of intratracheal infection and after 30 days, lower fungal load, when compared with Swiss mice. According to the set of experiments performed, our findings reveal that independently of mice strain, PMΦ is more reactive against *R. oryzae* in the first contact than AMΦ. In addition, increased PMΦ production of H_2_O_2_ at the end of disseminated infection is related to efficient fungal clearance observed in resistant (BALB/c). Our findings provide new evidence to understand the parasite-hosts relationship in mucormycosis.

## 1. Introduction

Mucormycosis is a devastating invasive fungal infection predominantly caused by *Rhizopus* species (1). It has emerged globally as a public health threat during the COVID-19 pandemic (2–4). The infection occurs via diverse routes such as inhalation, percutaneous, or ingestion of spores. Mucormycosis may have a wide variety of disease manifestations, with rhinocerebral and pulmonary diseases being the most common (5). The mortality rate varies from 46%-70% and up to 90% upon dissemination (6–8).

Risk factors for mucormycosis include diabetes mellitus, neutropenia, sustained immunosuppressive therapy, chronic prednisone use, and iron chelation therapy (7). During the COVID-19 pandemic, the use of immunomodulatory drugs (e.g., systemic corticosteroids and tocilizumab) and COVID-19-induced immune dysregulation increased the risk of mucormycosis (9,10). Cases of COVID-19–associated mucormycosis (CAM) have been reported worldwide (11–20), and have been mostly associated with underlying uncontrolled diabetes mellitus diseases. Nevertheless, cutaneous and soft-tissue mucormycosis has been observed in immunocompetent individuals (21,22), such as those induced by traumatic implantation of contaminated soil and water during tornadoes (23).

Considering the high rates of mortality and morbidity associated with this life-threatening disease (14), it has a prognosis and outcome that have not significantly improved over the last decades, mainly due to the difficulty in forming an early diagnosis and the limited activity of current antifungal agents against Mucorales (24,25). In addition, mucormycosis has an incompletely understood pathogenesis, particularly related to the parasite-hosts relationship.

Experimental models of mucormycosis are widely used for the evaluation of antifungal therapy and, in these studies, immunosuppression is used to induce fungal dissemination (26–29). Nevertheless, the effect of immunosuppressive drugs restricts the evaluation of the immunological mechanisms involved in fungal resistance. Recently, our group has focused on studies exploring *Rhyzopus*-host interplay using immunocompetent models, inducing both disseminated and pulmonary mucormycosis (30,31). Using BALB/c and Swiss mice, we determined a resistant and less resistant model of mucormycosis, respectively. While BALB/c mice has been shown better capacity in decreasing fungal load until 30 days of intravenous and intratracheal infection, Swiss mice has been shown be less responsive to *R. oryzae* infectious and consequently more prolonged viable fungal presence in internal organs (30,31). We and others (32,33) have highlighting the importance of evaluating immune response in immunocompetent mice to understand the mechanisms involved in the resistance to Mucorales agents, as well as, the mucormycosis pathogenesis in immunocompetent individuals (21,22).

Andrianaki and collaborators (2018) revealed the essential role of *Rhizopus*– macrophage interplay for pulmonary mucormycosis. Using an immunocompetent model, it was shown that the development of mucormycosis is related to the prolonged intracellular survival of the fungus inside of the macrophages (33). Considering the poor knowledge of macrophage activity during mucormycosis in the context of natural resistance; in the present study, we employed resistant and less resistant models to determine the reactive oxygen and nitrogen species production by *R. oryzae*-infected macrophages

## 2. Material and methods

### 2.1 Mice

Two-month-old female inbred BALB/c and outbred Swiss mice from the Animal House at the Laboratório de Imunopatologia Experimental (LIPE) of UNESP (Univ Estadual Paulista, Bauru, SP, Brazil) were divided into groups randomly. We provide food and water sterile *ad libitum*. This study was carried out in strict accordance with the recommendations in the Guide for the Care and Use of Laboratory Animals of the National Institutes of Health and Brazilian College of Animal Experimentation. The study was approved by an Institutional Animal Care and Use Committee (IACUC) (Protocol Number: 1608/46/01/2013-CEUA-FC) by Animal Experimentation Ethics Committee of the School of Sciences of Bauru, UNESP. All surgery was performed under ketamine/xylazine anesthesia, and all efforts were made to minimize suffering.

### 2.2 Fungal strains

*R. oryzae* (IAL 3796) was previously obtained from the fungal collection of Instituto Lauro de Souza Lima (ILSL), and submitted to species identification by Adolfo Lutz Institute (São Paulo, SP, Brazil). The fungi were maintained by monthly subculturing on Sabouraud dextrose agar (SDA) slants (Difco Laboratories, Detroit, Michigan, USA).

### 2.3 Experimental design

Mice were randomly separated into two main groups according to the route of inoculation: intravenous groups (*Rhi-*IV) and intratracheal groups (*Rhi-*IT). For the *Rhi-*IV groups, Swiss and BALB/c mice were inoculated within 3.0 × 10^4^ viable spores of *R. oryzae* in the caudal vein. For the *Rhi-*IT groups, Swiss and BALB/c mice were inoculated within 2.0 × 10^6^ viable spores of *R. oryzae* in the trachea. Next, groups of six *R. oryzae*-infected mice were evaluated on days 7 and 30 post-infection (p.i.). Non-infected groups were composed of BALB/c and Swiss mice inoculated with sterile saline solution (SSS) by intravenous or intratracheal route.

### 2.4 Fungal infection

Fungi were washed carefully with SSS, and the suspension was mixed twice for 10 s on a vortex-mixer and decanted off for 5 min. The supernatants were collected and washed twice. We determined fungal viability by cotton blue staining. In *Rhi-*IVgroups, a volume of 100 μL of fungal suspension (3 × 10^4^ spores of *R. oryzae*) was inoculated into the lateral tail vein. In *Rhi-*IT groups, mice were previously anesthetized ketamine and xylazine (80 and 10 mg/kg body weight, respectively) by intraperitoneal route. After the tracheal exposition, each mouse received a volume of 40 μL (2 ×10^6^ spores of *R. oryzae*) of the suspension. The incision was sutured with surgical thread, and the mice were kept in a warm place and observed for their recovery process.

### 2.5 Collection of the biological material

Mice were anesthetized with isoflurane then euthanized via CO_2_ asphyxiation. Peritoneal lavage (PL) and bronchoalveolar lavage (BAL) was performed in *Rhi-*IV and *Rhi-*IT groups, respectively, using cold and sterile phosphate-buffered saline (PBS). Fragments of the brain, liver, lung, spleen, and kidneys were collected and submitted to microbiological evaluation.

### 2.6 Recovery of viable fungi

Quantitative colony culture is a wide method for organ fungal burden determination in per gram tissue in which whole tissue is grinding into suspension before inoculating onto culture plates. However this methods causes non-viability in Mucorales (21), and to avoid damage to fragile hyphal and consequent false-negative results in our quantitative analysis (34), the fragment method was employed in our set of experiments, as previously reported (30,31,35). For this, ten fragments (2 × 2 mm) of the brain, liver, lung, spleen, and kidneys were cultured on Sabouraud agar plates at 25ºC for a maximum of seven days. We counted fragments of fungal growths and expressed the results as the frequency of *R. oryzae*-positive fragments per total of cultivated.

### 2.7 Macrophages cell culture

BAL and PL suspensions were centrifuged for 10 minutes at 410 *g*, and cells were resuspended in 1.0 mL of supplemented RPMI-1640 (Nutricell, Campinas, SP, Brazil). The supplemented RPMI-1640 contained: 10% heat-inactivated fetal calf serum (Nutricell), penicillin (100 UI mL^-1^), streptomycin (100 mg mL^-1^) (Sigma-Aldrich, St. Louis, MO, USA), and amphotericin B (0.25 μg mL^-1^) (Sigma-Aldrich). Cell concentration was adjusted to 1.0 × 10^5^ mononuclear phagocytes mL^-1^ as judged by the uptake of 0.02% neutral red (Sigma) and confirmed by expression of F4/80 by Fluorescence-Activated Cell Sorting (FACS). Cells were plated into 96-well flat-bottomed microtiter plates (Greiner BioOne, Frickenhausen, Germany), and incubated for two hours at 37°C and 5% CO_2_ in a humidified chamber to allow cells to adhere and spread. Non-adherent cells were removed by washing the wells three times with RPMI, and we used the remaining adherent cells (>95% mononuclear phagocytes as assessed by morphological examination) for experiments. The adherent cells were cultured at 37°C and 5% CO_2_ in supplemented RPMI-1640 with or without heat-killed spores of *R. oryzae* (*R. oryzae*-Ag) (1 spore: 1 cell). As an internal control for macrophage activity, cells were cultured with 10 μg ml-1 lipopolysaccharide (Sigma-Aldrich). After 24 hours, cell-free supernatants were harvested and stored at −80°C for cytokine analysis.

### 2.8 Production of hydrogen peroxide (H_2_O_2_)

The production of H_2_O_2_ was estimated according Russo et al. (1989). Briefly, adhered mononuclear phagocytes obtained as described before were maintained in RPMI-1640 culture medium at 37°C and 5% CO_2_ for 24 h. At the end of the cell culture period, we removed the supernatants from the wells, added the phenol red solution containing dextrose (Sigma), phenol red (Sigma), and horseradish peroxidase type II (Sigma), and incubated the plate at 37°C in 5% CO_2_ for one hour. Reaction was stopped by adding 1 N NaOH. The H_2_O_2_ concentration was determined using a colorimetric microreader (ELx 800; BioTek Instruments Inc., Winooski, VE, USA)(36).

### 2.9 Detection of levels of nitric oxide

**(NO)**. The production of NO was estimated following the Griess’ method (Green et al., 1982). Production of nitrite, a stable end product of NO, was measured in the cell-free supernatants of adhered mononuclear phagocytes cultured. Briefly, a volume of 0.1 mL of cell-free supernatant with an equal volume of Griess reagent was incubated for 10 minutes at room temperature. Griess reagent was prepared using: 1% sulfanilamide (Synth, Diadema, SP, Brazil), 0.1% naphthalene diamine dihydrochloride (Sigma), and 2.5% H3PO4. The nitrite accumulation was quantified using a colorimetric microreader (ELx 800; BioTek Instruments Inc., Winooski, VE, USA). Nitrite concentration was determined using sodium nitrite (Sigma) diluted in RPMI-1640 medium as a standard (36).

### 2.10 Dosage of TNF-α and IL-10

IL-10 and TNF-α levels were measured in the cell-free supernatants of cell cultures using a cytokine Duo-Set Kit (R&D Systems, Minneapolis, MI, USA), according the manufacturer’s instructions.

### 2.11 Statistical analyses

Normality tests of data was determined by Shapiro–Wilk’s test. For comparing two independent samples *t*-test was performed. For multiple comparisons, ANOVA with the Tukey post-test was used. Pearson’s correlation coefficient was determined to measure the statistical association between two continuous variables. All statistical tests were performed using GraphPad Prism version 5.0 for Windows (GraphPad Software, San Diego, CA). *P*-value of 5% or lower was considered statistically significant (Zar, 2010).

## 3. Results

### 3.1 Production of reactive oxygen reactive species and IL-10 by heat-killed *R. oryzae*-infected peritoneal macrophages is dependent of mouse genetic background

First, we evaluated the *in vitro* response of peritoneal (PMΦ) and alveolar (AMΦ) macrophages obtained from non-infected BALB/c and Swiss mice challenged with heat-killed *R. oryzae* (*R*.*oryzae*-Ag). To evaluate the PMΦ and AMΦ response we measured reactive oxygen and nitrogen species (H_2_O_2_ and NO) in the cultures. We also measured the levels of TNF-α and IL-10 to validate our findings.

In the presence of heat-killed spores of *R. oryzae*, PMΦ from the two mouse strains showed decreased production of the NO (Fig 1A) and increased the TNF-α production (Fig 1C). Interestedly, *R. oryzae*-infected PMΦ from BALB/c showed increased production of H_2_O_2_ and IL-10. (Fig 1B, Fig 1D).

**Fig 1.**
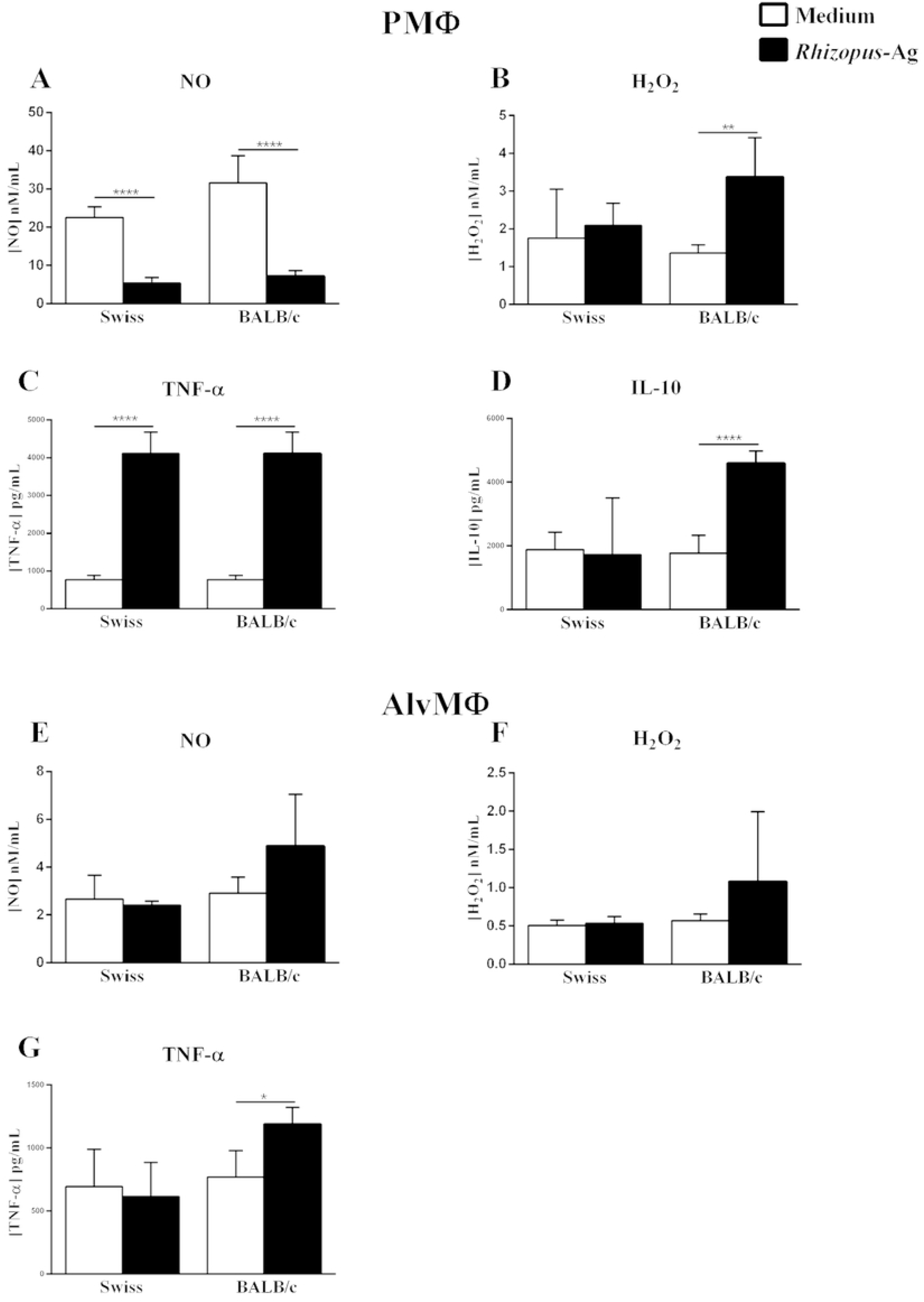
Peritoneal and alveolar macrophages from naive Swiss and BALB/c mice react differently in front of heat-killed *R. oryzae*. (A) NO, (B) H2O2, (C) TNF-α, (D) IL-10 levels in cell-free supernatant of peritoneal macrophages, and (E) NO (F) H2O2, (G) TNF-α levels in cell-free supernatant of alveolar macrophages of non-infected Swiss, and BALB/c mice co-cultured or not with heat-killed spores of R. oryzae. Student’s t-test; n=5-7; *p < 0.05, ** p< 0.01, ***p< 0.001). NO: nitric oxide; H2O2: hydrogen peroxide; TNF-α: tumor necrosis factor-alpha; IL-10: interleukin 10; PMΦ: peritoneal macrophages; AlvMΦ: alveolar macrophages.

In general, AMΦ from both Swiss and BALB/c mice showed similar production of NO and H_2_O_2_ (Fig 1E, Fig 1F). A slight increased production of TNF-α by *R. oryzae*-infected AMΦ obtained from BALB/c was observed. In this set of experiments, no detectable production of IL-10 was observed.

### 3.2 Enhanced production of H_2_O_2_ by peritoneal macrophages was associated with better clearance of *R. oryzae* in a model of disseminated mucormycosis

In the present study, BALB/c and Swiss mice strains showed more and less resistance, respectively, against *in vivo R. oryzae* infection, as previously reported (30,31). The course of infection was characterized by similar fungal load on day 7 for both mouse strain, followed by a decreased fungal load on day 30, that was more expressive in BALB/c mice (Fig 2A).

**Fig 2.**
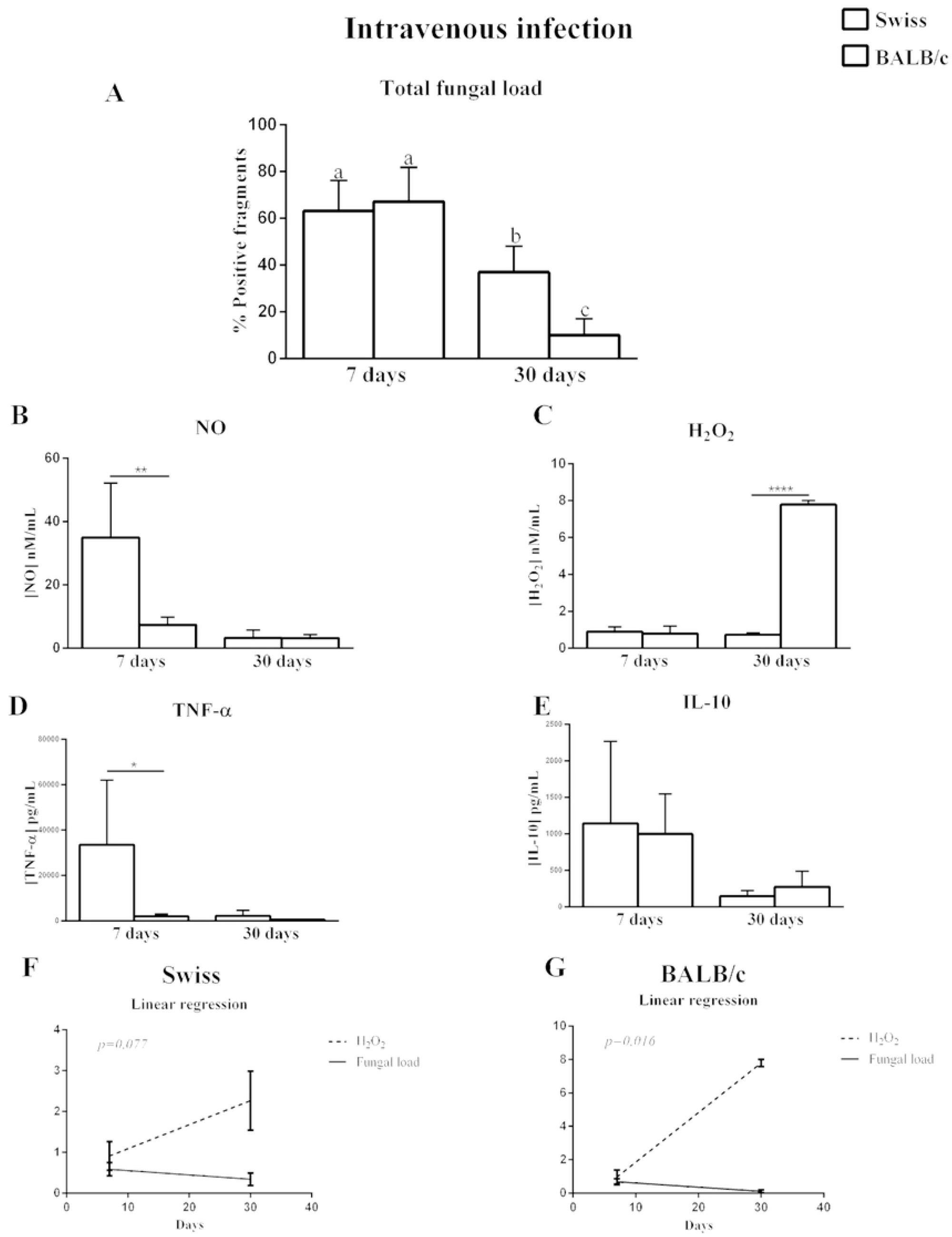
Increased PMΦ production of H_2_O_2_ at the end of disseminated infection is related to efficient fungal clearance in BALB/c mice. (A) Total fungal load (B) NO, (C) H_2_O_2_, (D) TNF-α, and (E) IL-10 levels in cell-free supernatant of peritoneal macrophages from Swiss or BALB/c mice co-cultured with heat-killed spores of *R. oryzae*. Linear regression analysis between H_2_O_2_ levels and total fungal load in BALB/c (F) and Swiss (G) mice. The infected group was composed of mice inoculated intravenously with 3 × 104 spores of *R. oryzae* and evaluated after 7 and 30 days. Any significant differences relative to infected samples compared to different times post infection (letters) and to different strains (*) are indicated. (Student’s t-test; n=5-7; *p < 0.05, ** p< 0.01, ***p< 0.001). NO: nitric oxide; H_2_O_2_: hydrogen peroxide; TNF-α: tumor necrosis factor alpha; IL-10: interleukin 10.

In order to explore the results of *in vitro R. oryzae-*infected macrophages experiments, we hypothesized that the production of reactive species of oxygen and nitrogen by macrophages could be involved in the differential response observed between these two mice strains.

To assess our hypothesis, we first evaluated the specific activity of PMΦ derived from BALB/c and Swiss mice that were intravenously infected with *R. oryzae*. For this, on days 7 and 30 p.i., PMΦ were recovered from the peritoneal cavity and challenged with heat-killed spore of *R. oryzae*.

On day 7, PMΦ from Swiss mice showed higher production of NO (Fig 2B) and TNF-α (Fig 2D) than PMΦ from BALB/c mice. On the other hand, on day 30, more resistant BALB/c mice showed higher levels of H_2_O_2_ (Fig 2C). No differences in the production of NO, TNF-α, and IL-10 were observed between both mouse strains (Fig 2B, Fig 2D, Fig 2E).

Next, we performed a correlation test between production of H_2_O_2_ and fungal load. An inverse correlation was observed between the production of H_2_O_2_ and total fungal load in the BALB/c mice (Fig 2G), but not in the Swiss mice (Fig 2F).

### 3.3 Enhanced initial pro-inflammatory response and NO production by alveolar macrophages of BALB/c mice seems to better control the pulmonary mucormycosis

BALB/c and Swiss mice were intratracheally infected with *R. oryzae* to evaluate the tissue-specific response. As observed in intravenous infection, both BALB/c and Swiss mice showed a decrease in viable fungal load after 30 days p.i., and BALB/c mice showed better fungal clearance than Swiss mice (Fig 3A).

**Fig 3.**
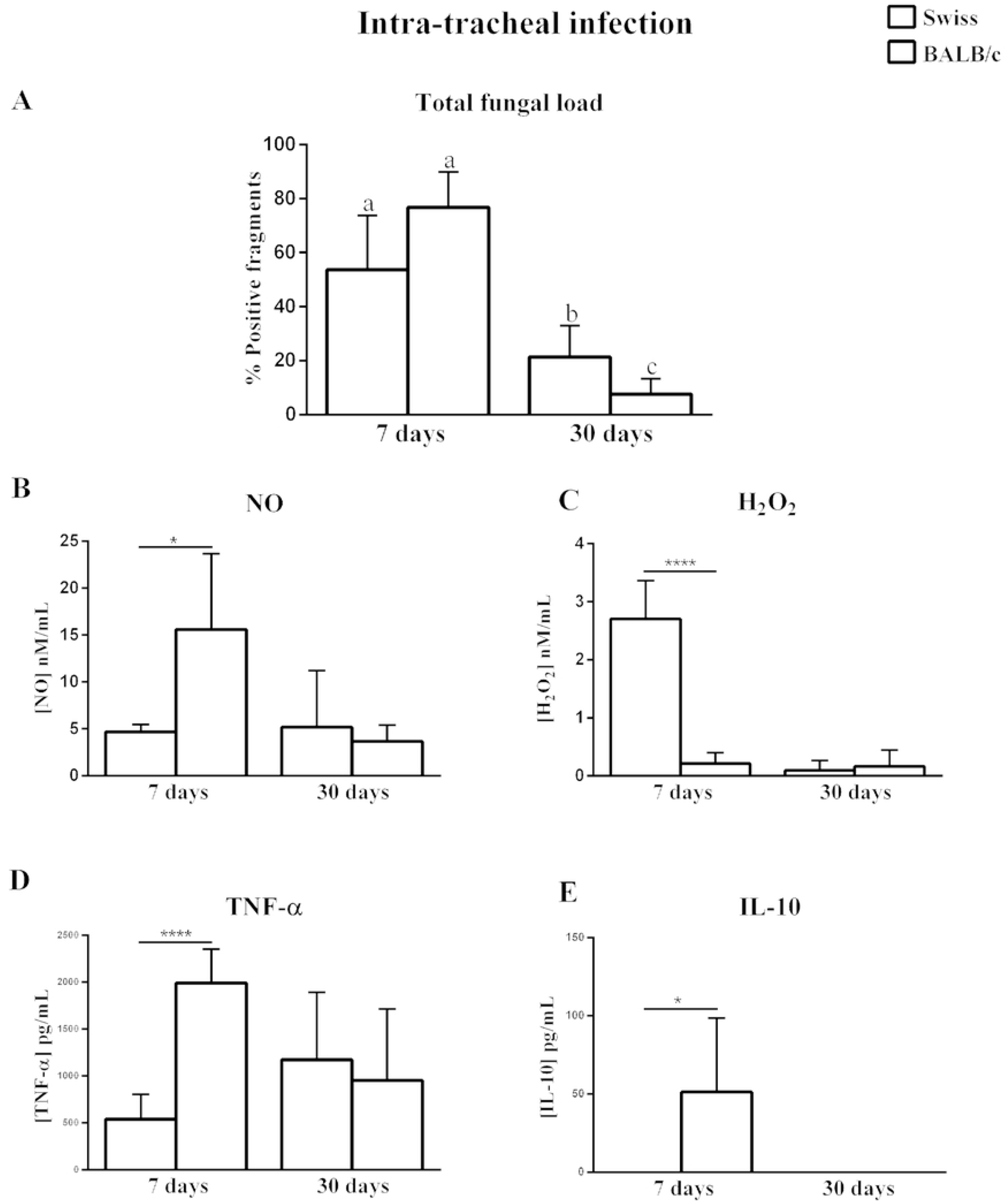
AMΦ from BALB/c mice showed higher production of NO, TNF-α, and IL-10 than AMΦ from Swiss mice after 7 days of *R. oryzae* pulmonary infection. (A) Total fungal load, (B) NO, (C) H_2_O_2,_ (D) TNF-α, and (E) IL-10, levels in cell-free supernatant of alveolar macrophages from Swiss or BALB/c mice co-cultured with heat-killed spores of *R. oryzae*. The infected group was composed of mice inoculated intratracheally with 2 ×106 spores of *R. oryzae* and evaluated after 7 and 30 days. Any significant differences relative to infected samples compared to different times post infection (letters) and to different strains (*) are indicated. (Student’s t-test; n=5-7; *p < 0.05, ** p< 0.01, ***p< 0.001). NO: nitric oxide; H_2_O_2_: hydrogen peroxide; TNF-α: tumor necrosis factor alpha; IL-10: interleukin 10.

On day 7, AMΦ from BALB/c mice showed higher production of NO (Fig. 3B), TNF-α (Fig 3D), and IL-10 (Fig 3E) than AMΦ from Swiss mice. Lower production of H_2_O_2_ by AMΦ from BALB/c mice was observed (Fig 3C).

On day 30, no differences in the production of NO, H_2_O_2_, TNF-α, and IL-10 by AlvMΦ were observed (Fig 3B, Fig 3C, Fig 3D, Fig 3E).

## 4. Discussion

Here we use the different outcomes observed in experimental mucormycosis between the immunocompetent BALB/c and Swiss mice to highlight the protective pattern of macrophages responses against *R. oryzae*. We found that macrophages from different microenvironments showed different responses against *R. oryzae* not only *in vitro* but also during *in vivo* infection. Also, the better outcome was associated with higher H_2_O_2_ produced by PM, and higher initial NO, TNF-α, and IL-10 produced by AMΦ during experimental disseminated and pulmonary mucormycosis, respectively. In addition, the genetic background interferes in the macrophage response, so, regardless of the microenvironment, the BALB/c’s macrophage is more effective in eliminating the fungus.

Inbred mouse strains vary widely in their degree of innate susceptibility to systemic fungal infections (37–41). BALB/c and Swiss mice are resistant and susceptible strains, respectively, in experimental infections caused by *Cryptococcus neoformans* (Rhodes et al., 1980), *Candida albicans* (38,42), and *Paracoccidioides brasiliensis* (41). It was identified that the genetic target C5-deficiency (Hc0 allele, hemolytic complement) modulates the host’s initial response and causes susceptibility and ineffective inflammatory response against fungal infections (38,39). Active as an anaphylatoxin, the C5 molecule is a power chemotactic factor for polymorphonuclear leukocytes. Lack of C5 would block complement, thereby decreasing the opsonization and recruitment of phagocytic cells (39). Although the higher susceptibility of Swiss mice to fungal infections may be explained by a deficiency in the Hc0 allele (41–43), our results suggest that other genetic targets may be involved in the less responsiveness of macrophages from non-infected Swiss mice to heat-killed *R. oryzae*.

According to clinical and experimental data, individuals who lack phagocytes or present impaired phagocytic functions show a higher risk of developing mucormycosis (6,44,45). We observed in the *in vitro* part of this study that peritoneal macrophages from the most resistant strain (BALB/c) showed a potent inflammatory response mediated by H_2_O_2_ and TNF-α. (46) in the first contact with *R. oryzae* antigen. A previous *in vitro* study showed that the inactivated Mucorales cells can stimulate high levels of TNF-α, IL-1β, IL-6, I-L8, GM-CSF, and MCP-1 production by humans cells of different immune subsets, with paralleled upregulation of transcriptional activity of IL-1β and TNF-α (47,48). Since the oxidative microbicidal production by neutrophils and monocytes is related to *R. oryzae* hyphae damage and killing (49,50), our results indicate an initial proinflammatory response against *R. oryzae* can lead to a better outcome in experimental mucormycosis.

Unexpectedly, in the first contact with *R. oryzae* antigen peritoneal macrophages, in general, decreased the production of NO. Similar to our results, it was demonstrated that *Aspergillus nidulans* melanin inhibited the NO production by lipopolysaccharide (LPS)-stimulated peritoneal macrophages, accompanied by a slight stimulatory effect on TNF-α production (51). Melanin is a type of pigment in the fungal cell wall of black molds and has been shown to block the effects of hydrolytic enzymes on the cell wall (52). Considering the importance of this finding, we suggest that the immunomodulatory effects induced by the *R. oryzae* cell wall on peritoneal macrophages need to be better investigated.

Differently from peritoneal macrophages, alveolar macrophages from the BALB/c strain produced higher NO and TNF-α levels than less resistant strain, during the *in vivo* model of pulmonary mucormycosis. It is known that pulmonary activated macrophages are the major defense against fungal invasion (53). In contact with macrophages receptors, TNF-α provides signals that lead to an induction of antimicrobial activity. This activity depends on NO synthase activation and NO production (54). Some studies are demonstrating that this process is essential to fungal death. In paracoccidioidomycosis (PCM), TNF-a induces *P. brasiliensis* killing by H_2_O_2_ and NO release (55). In cryptococcosis, TNF-α significantly promoted macrophage NO production and anticryptococcal activity(56). In addition, TNF-α enhances pulmonary alveolar macrophage phagocytosis and oxygen production during initial contact with *A. fumigatus* (57). It is known that TNF-α contributes to the influx and activation of neutrophils and mononuclear cells in the lungs during the filamentous fungal challenge (58). In mucormycosis, the TNFα signaling may have a protective response since patients treated with a tumor necrosis factor inhibitor have a higher risk to developed disseminated mucormycosis(59,60).

Although the association of NO with the killing of yeast cells supports the importance of this metabolite in host protection (61), other studies have been demonstrating that the overproduction of this metabolite is associated with susceptibility in experimental PCM (62,63). Nascimento, et. al (2002) observed that high levels of NO induce T cell immunosuppression during *P. brasiliensis* infection (Nascimento et al., 2002). These reports suggest that the protective or deleterious NO role *in vivo* depends on the balance between its fungicidal and immunosuppressive properties. Our data demonstrate that after intravenous *R. oryzae* infection, stronger and early production of NO and TNF-a by peritoneal macrophages from Swiss mice does not result in late better fungal clearance. On the other hand, after a pulmonary *R. oryzae* infection, higher and early production of NO and TNF-a by alveolar macrophages from BALB/c mice was accompanied by higher levels of IL-10, resulting in later better control of pulmonary mucormycosis. Suggesting that in BALB/c mice, an initial balanced immune response is essential for better infection control.

IL-10 is an anti-inflammatory cytokine that can impede pathogen clearance. But it also can ameliorate immunopathology (64). The balance between pro-and anti-inflammatory cytokines is crucial for the host’s defense against *A. fumigatus*. An *in vitro* study with mononuclear cells (MNC) stimulated with *A. fumigatus* showed that hyphae of this fungi induced the release of IL-10 by MNC, and this process was dependent on endogenous IL-1 (65). Whereas inflammatory cytokines such as IFN-γ, TNF-α, and IL-18 activate monocytes and neutrophils to ingest and kill *A. fumigatus* conidia and hyphae, the subsequent release of the anti-inflammatory cytokine IL-10, are responsible for down-regulating the potential deleterious overstimulation induced by the inflammatory mediators (66).

In general, one of the biological effects of IL-1 is an increased synthesis of IL-10 (67). Netea et al., (2004) showed that *Candida* stimulated IL-1 release by cells, and secondarily IL-10 release (68). In Mucormycosis, although high production of IL-10 by Mucorales-specific T cells from patients was related to susceptibility in the late phase of the disease (69), our results showed a correlation between higher IL-10 production by alveolar macrophages in the initial moments of *R. oryzae* infection and better outcome in experimental pulmonary mucormycosis. We suggest that high levels of IL-10 were a biological effect to balance the pro-inflammatory response mediated by high levels of TNF-α and NO. A fundamental step for an efficient immune response.

While alveolar macrophages from a more resistant strain showed an initial higher TNF-α, NO, and IL-10 response to a pulmonary infection, the peritoneal macrophages from the same strain showed a late and strong response mediated by H_2_O_2_ after the intravenous *R. oryzae* infection. It is important to note that after infection by the intravenous route, the decrease in viable *R. oryzae* was inversely proportional to the release of H_2_O_2_ by peritoneal macrophages from a more resistant strain of mice. It indicates the relation of this mediator with protection in Mucormycosis. This result agrees with the study done by Andrianaki et al. (33), where they prove the susceptibility of *A. fumigatus* and *R. oryzae* conidia to oxidative damage induced by H_2_O_2_. In this context, an important detail was that the BALB/c strain only showed a more efficient fungal clearance after 30 days of infection. We suggest that the adaptative immune response potentiated the activity of the macrophages evaluated in this study.

Macrophages are the first immune cells to interact with invasive pathogens. Depending on macrophage’s activity, they can prime the adaptive immune response, which is far more aggressive and specific to pathogens (70). Generally, high levels of ROS production by macrophages are linked to intracellular events, including the activation of transcription factors such as NFκB (71), and induction of mitogenesis (72). In addition, *in vitro*, and *in vivo* studies already showed that *R. oryzae* could trigger a Th-17 response mediated by high amounts of IL-23 by dendritic cells (73), and also the association of high levels of IL-17 and IFN-γ with better *R. oryzae* elimination by immunocompetent BALB/c and C57BL/6 mice (30,31). Considering the studies above and the evidence found here, we hypothesized that adaptive immune responses developed by BALB/c mice were essential to enhance the fungicidal activity of peritoneal macrophages and efficiently kill *R. oryzae*.

The absence of additional experiments to explore the intracellular mechanisms mediating the differences in ROS production and to confirm the adaptative immune response related to a more efficient macrophages response front of *R. oryzae*-Ag was the main limitation of the present study, and we suggest that more studies should be done to clarify the results observed here.

In summary, our findings reveal that independently of mice strain, PMΦ is more reactive against *R. oryzae* in the first contact than AMΦ. In addition, increased PMΦ production of H_2_O_2_ at the end of disseminated infection is related to efficient fungal clearance observed in resistant (BALB/c). Our findings provide new evidence to understand the parasite-hosts relationship in mucormycosis.

## Striking image. Overview of *in vitro* and *in vivo* macrophages response in BALB/c and Swiss mice infected with *R. oryzae* via different routes of infection

Macrophage activity of BALB/c and Swiss mice were evaluated pre-and-post intravenous or intratracheal *R. oryzae* infection. NO, H_2_O_2_, TNF-α, and IL-10 concentrations were evaluated in macrophages co-cultured or not with heat-killed spores of *R. oryzae*. Infected mice were intravenously or intratracheally inoculated with 3 × 10^4^ and 2 ×10^6^ spores of *R. oryzae* respectively, and evaluated after 7 and 30 days. Summarized results of the fungal load on day 30 are also shown. NO: nitric oxide; H_2_O_2_: hydrogen peroxide; TNF-α: tumor necrosis factor alpha; IL-10: interleukin 10; PMΦ: peritoneal macrophages; AlvMΦ: alveolar macrophages. Image by Débora Almeida-Donanzam via Inkscape

